# A New Method for Optimal Placement of Tumor Treating Fields Electrodes

**DOI:** 10.1101/2025.11.19.689304

**Authors:** Konstantin Weise, Nikola Mikic, Fang Cao, Eric T. Wong, Thomas R. Knösche, Axel Thielscher, Anders Rosendal Korshøj

**Author notes:** **Corresponding Authors:** Konstantin Weise, Dr.-Ing., Address: Max Planck Institute for Human Cognitive and Brain Sciences, Stephanstr. 1a, 04103 Leipzig, Germany, Anders Rosendal Korshøj, MD, PhD, Address: Aarhus University Hospital, Department of Neurosurgery, Palle Juul-Jensen’s Boulevard 165, J604, 8200 Aarhus N, Denmark. contributed equally (Axel Thielscher, Anders Rosendal Korshøj). **List of unpublished papers cited:** Mikic NM, Lukacova S, Skjøth-Rasmussen J, et al. (2025) Dose-enhanced vs standard TTFields for first recurrence of glioblastoma. A randomized clinical trial (resubmitted to Neuro-Oncology in October, 2025, Neuro-Oncology Advances NOA-D-25-00312.).

## Abstract

**Overview:** Tumor Treating Fields (TTFields) provide a non-invasive treatment option for newly diagnosed glioblastoma. While optimization of electrode placement is important to increase treatment efficacy, clinical therapy planning is done using an undisclosed and proprietary software (NovoTAL®), which is clinically unvalidated. This study investigates a new computational approach for optimizing TTFields electrode placement and is compared to the current clinical standard.

**Methods:** We developed a new computational pipeline integrating patient-specific anatomical data to optimize electrode configurations in five representative glioblastoma cases with diverse tumor locations and sizes. Two optimization strategies were employed: one maximizing electric field intensity at the tumor, and another enhancing coverage of the adjacent brain while maintaining sufficient tumor intensity. Results were compared to electrode placements generated by NovoTAL®. Additional simulations with artificial tumors assessed the effects of tumor size and location.

**Results:** Optimized electrode placements improved electric field intensity in tumors by 18%–34% compared to the clinical standard. Coverage-weighted optimizations provided broader field coverage without significantly compromising tumor intensity. Smaller or surface-adjacent tumors benefited most from optimization, achieving precise targeting and enhanced coverage. Extensive randomized placement analyses highlighted the superior performance of the optimized configurations. Analysis of artificial models showed consistent improvements across varying tumor locations and sizes.

**Conclusion:** Personalized optimization of TTFields electrode placement significantly improves electric field targeting of tumors and adjacent brain regions. This approach outperforms standardized planning software and clinical practices and supports future development of adaptive, automated strategies for individualized TTFields therapy in glioblastoma.

**Keypoints:**

- Optimized TTFields array placement enhanced field intensity by 18–34% vs. clinical standard.
- Optimized TTFields planning improved field coverage in tumor-adjacent regions.
- Optimized TTFields planning outperformed standard methods and random placement.

**Importance of the study:** TTFields are increasingly used as adjuvant therapy for glioblastoma. However, current individualized treatment planning relies on proprietary, undisclosed, and clinically unvalidated software, limiting transparency, optimization, and innovation in the field. This study introduces an individualized, semi-automated, and open-source method for optimal electrode placement based on standard MRI data, addressing a critical need for validated and adaptable planning tools. Our approach increased field intensity in tumors by 18–34% and achieved broader coverage compared to the standard clinical tool (NovoTAL®), without compromising therapeutic strength. Notably, the method also consistently outperformed extensive random electrode placements across diverse tumor types and sizes, highlighting its robustness and ability to achieve true optimal configurations. Open-source availability enhances reproducibility and clinical translation, representing a significant step toward more effective, individualized TTFields therapy. This advancement has the potential to improve outcomes for glioblastoma patients and underscores the importance of technology validation in neuro-oncology.

## Introduction

The management of glioblastoma continues to pose significant challenges, motivating the development of new therapeutic strategies. One such approach is Tumor Treating Fields (TTFields), a non-invasive treatment that employs alternating electric fields (E-fields) to disrupt cancer cell proliferation. First introduced by Kirson et al. in early clinical trials^1^, TTFields have demonstrated efficacy against solid cancers^2^, including newly diagnosed glioblastoma^3^, and non-small cell lung cancer^4^. The electric fields are thought to exert forces on charged or polarizable molecules within dividing cells, disrupting critical structures, including the mitotic spindle and membrane proteins, ultimately causing cancer cell death^5,6^.

The intensity of TTFields is believed to constitute an important dose-metric correlating with the efficacy of the treatment^7^. Specifically, preclinical studies have demonstrated enhanced anti-neoplastic effects at increased field intensities across diverse cancer types^1,8^. Moreover, post hoc analyses of data from the EF-14 trial^3^ demonstrate an association between high field intensities in the tumor, longer overall survival^9^, and a reduced risk of local recurrence^10^. This dose-response correlation has prompted the development of the proprietary software NovoTAL® (Novocure, Ltd.), which is used together with the TTFields technology Optune® (Novocure, Ltd.), to prepare an optimized electrode layout for each patient. This concept compares to the principle of dose-planning in radiation oncology and is based on the fact that the placement of TTFields electrodes on the patient’s scalp directly influences the distribution and intensity of the electric fields, and hence expectedly impacts the therapeutic outcome. To achieve maximum clinical benefit of TTFields across cancer types, methods are needed to ensure the optimal electrical energy deposition in the regions of interest^11^.

The main objective of electrode planning, therefore, lies in determining individual electrode configurations that ensure optimum field intensity in the tumor. For glioblastoma, which infiltrates the adjacent brain tissue diffusely, maintaining high field intensities at tumor sites and adjacent regions bearing microscopic disease in mind is also important.

Currently, however, NovoTAL® is the only clinical tool available for TTFields electrode array planning. Its workflow is based on the morphometric inputs from structural MR images of the patient’s head (Fig. S1 in *Supplemental Material*). However, the underlying algorithm is not based on individual electric field simulations and is generally unknown. Moreover, its output is restricted to a choice between a few predefined and standardized layouts, raising questions about its accuracy and performance. The method’s superiority in terms of field optimization has not been validated against other optimization methods, and its clinical impact remains unexplored. Collectively, these factors represent important caveats to be addressed.

Recent advances in electric-field modelling of Tumor Treating Fields (TTFields) have established a detailed, multiscale understanding of how alternating electric fields interact with tissue and cellular structures. As summarized by Wenger and colleagues ^6,12^, state-of-the-art approaches employ volume conductor models solved using the finite element method under the electro-quasistatic approximation, enabling simulation of electric field distributions within patient-specific head models. This allows individualized electric-field estimates that reflect anatomical and biophysical variability across patients. Korshøj and colleagues^13,14^ extended these methods to analyze how tumor position, conductivity distribution, and skull geometry influence field strength, and how targeted craniectomy or optimized electrode placement can enhance local TTFields intensity. Furthermore, related methodological developments by Thielscher and colleagues in realistic head modelling for noninvasive brain stimulation^15^ have directly informed TTFields dosimetry pipelines.

In a recent study, we proposed a generalized optimization approach based on computational modeling and simulation of the electric field distributions to optimize electrode placement across diverse technologies, including Transcranial Electrical Stimulation (TES), Temporal Interference Stimulation (TIS), Electroconvulsive Therapy (ECT), and TTFields^16^. These simulations utilize advanced algorithms and finite element analysis to optimize the electric fields within complex anatomical structures to find optimal electrode arrangements by allowing free movement of the electrode arrays. By modeling patient-specific anatomical features based on standardized MRI data, it is possible to tailor electrode arrangements to ensure optimal therapeutic efficacy, such as maximal TTFields intensity within the tumor volume to enhance cytotoxic effects^17^.

In this study, we applied the electrode optimization methodology of Weise et al.^16^ to TTFields, illustrated in Fig. 1a, and evaluated its potential compared to the current clinical standard. For this purpose, we selected five representative patients with different tumor locations and sizes, and we compared the optimization method of Weise et al.^16^ to the electrode positions using the NovoTAL® planning software, which corresponds to the current clinical standard. We then juxtaposed the resulting electric fields in the tumor region from either method. Furthermore, we tested the method against extensive randomized array positioning to document robustness and true optimization across diverse tumor types and locations.

**Fig. 1:**
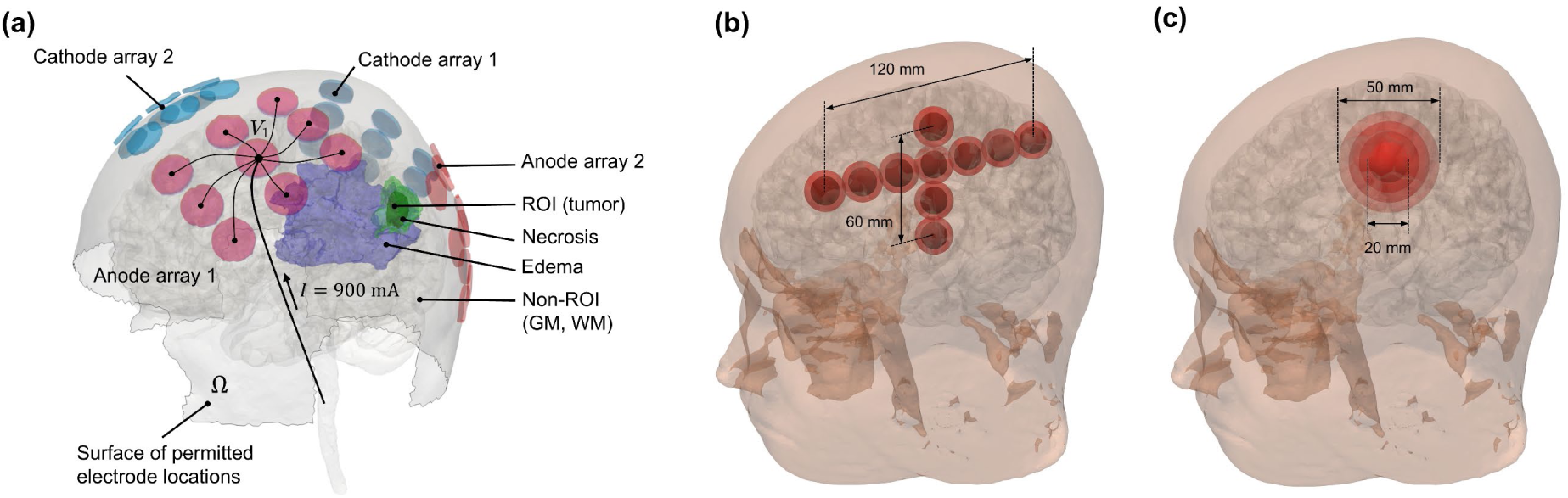
(a) Visualization of the TTFields electrode arrays on the patient’s head, together with the region-of-interest (ROI) containing the tumor and the “Non-ROI” containing the rest of the brain. The skin surface is restricted to locations where the electrode arrays can be placed, i.e. not over the eyes and ears of the patients; (b) modified *ernie* head model with different artificial tumor locations in superior-inferior (S-I) and anterior-posterior (A-P) direction and (c) tumor size to systematically study the impact of electrode optimization compared to current standard treatment strategies using TTFields.

## Methods

The optimization pipeline used to determine the optimal electrode placements for TTFields is implemented in SimNIBS v4.5^18^. All head models used in this study were created with T1 and T2 images using the CHARM pipeline^19^.

We studied real application scenarios on five patients representing a realistic variety of tumor morphologies and locations. Tissue volumes of edema, necrosis, and residual tumor were manually segmented using MRI patient data based on a standardized brain tumor protocol for MRI^20^. Following the Declaration of Helsinki, the study was reviewed and approved by the Central Denmark Region Committee for Health Research Ethics (68928/1-10-72-214-19) and the Danish Data Protection Authority. Written consent was obtained from the trial participants for using non-identifiable imaging for future in silico studies.

The electric conductivities for the pathological tissues were *σ_res.tumor_* = 0.24 S/m, *σ_necrosis_* = 1.0 S/m, and *σ_edema_* = 1.0 S/m ^13,14,21^. The electrical conductivities of the remaining tissues are given in Supplementary Material Table S1.

To systematically investigate the influence of tumor size and location, artificial tumors were added to the *ernie* head model, which is part of the example dataset of SimNIBS (https://simnibs.github.io/simnibs/build/html/dataset.html). The artificial tumor had a diameter of 20 mm and contained a necrotic inner part with a diameter of 14 mm. Its location was varied in longitudinal (superior to inferior, S-I) and sagittal (anterior to posterior, A-P) direction as shown in Fig. 1b. Additionally, the diameter of the tumor was expanded from 20 mm to 50 mm (Fig. 1c).

For each patient and the *ernie* head model, the required head and tumor measures were extracted from the MRI datasets and transferred to the planning software of NovoTAL® to determine the electrode positions of the current clinical standard. The software provides the suggested positions and orientations of the electrode arrays in the form of a graphical output using a general head template.

The electrode configurations were then manually applied to the computational head models to calculate the associated electric field distributions using SimNIBS v4.5. In addition, the electrode positions were determined using the optimization algorithm of SimNIBS v4.5 for comparison^16^. The algorithm determines the electrode array positions on the head that maximize a user-defined goal function. Two different goal function definitions were tested. In the first optimization approach (termed *intensity-optimized* in the following), the goal was to determine an electrode configuration that maximizes the average electric field strength in the tumor region. The second optimization approach (termed *coverage-optimized*) strived to maximize the spread of the electric field in the brain while also ensuring a field strength of 100 V/m in the tumor region, which is the dose for inhibiting tumor growth^1,8^. The second approach is motivated to sustain high field intensities throughout the rest of the brain to also target diffusely infiltrating cancer cells. In this context, the goal function is defined to maximize sensitivity while minimizing specificity. By defining the tumor and the rest of the brain as separate ROIs, the sensitivity and specificity can be optimized, given the chosen threshold of 100 V/m, by allowing the electrode arrays to move freely over the head surface. Details about the implementation of the optimization algorithm are given in Weise et al.^16^. Lastly, the final electric field distributions of the different optimizations were compared to the clinical standard using the NovoTAL® planning software.

## Results

Figure 2 presents a comparison of the current clinical standard with the optimized TTFields configurations for the five patients. The corresponding average electric fields and coverage scores are provided in Table 1. Across all cases, the individual optimizations consistently yielded higher average electric field strengths within the tumor region, ranging from 18% to 34%, relative to the current clinical standard. This improvement was observed for both intensity-optimized and coverage-optimized configurations. Depending on the chosen optimization criterion, either the intensity or the field coverage was enhanced, albeit with a slight trade-off in the other metric. For example, in the patient with a tumor in the left occipital lobe (first row in Fig. 2), switching from an intensity-optimized to a coverage-optimized electrode setup resulted in a trade-off: the intensity increase decreased from +22.8% to +13.8% relative to the clinical standard, while the coverage improved from −6.9% to −13.4%. The average trade-off over all patients was 8.7% in intensity and 10.3% in coverage score.

**Fig. 2:**
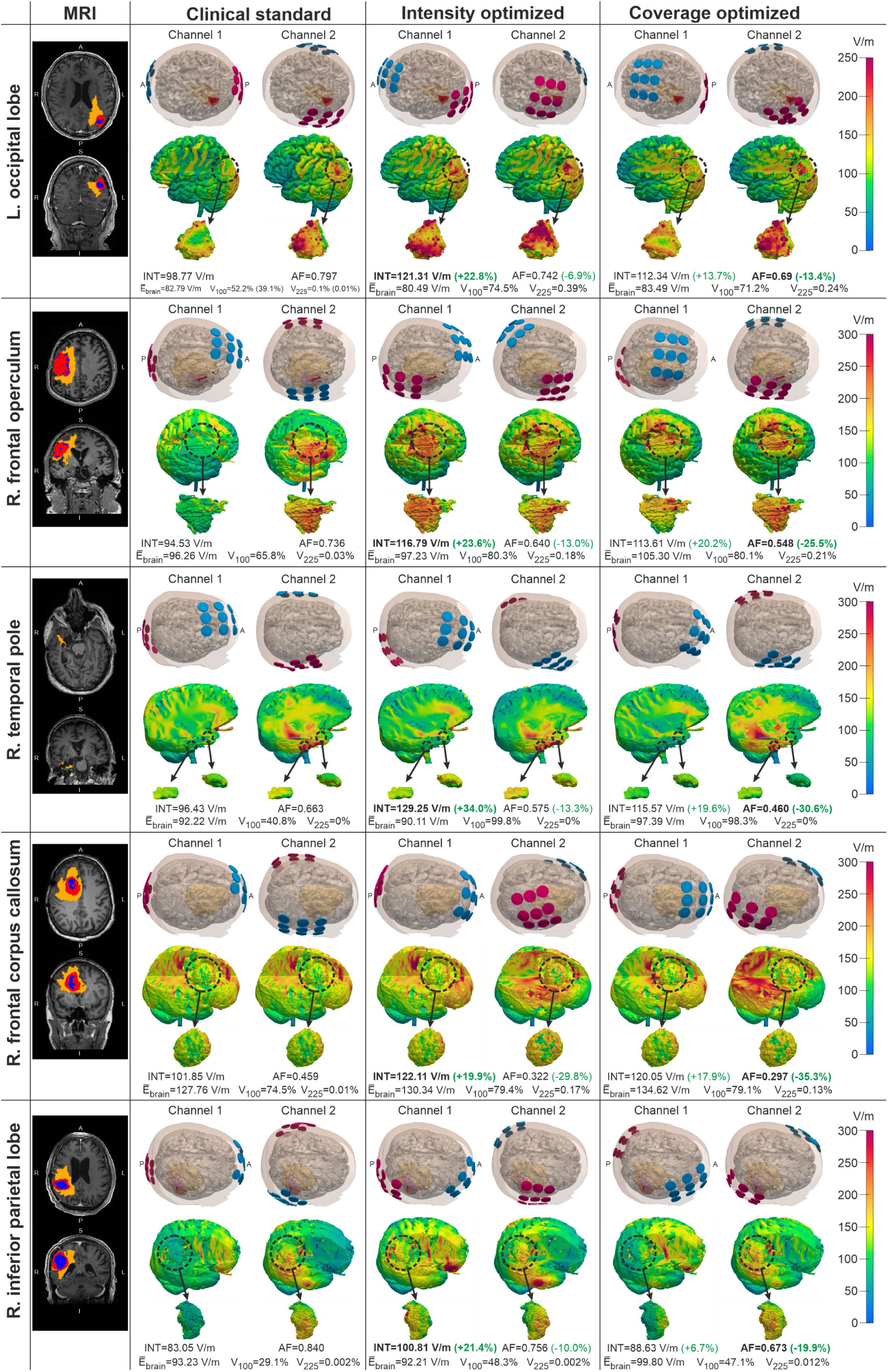
Electric field distributions created from both stimulation channels. The rows show the different patients. The first column shows axial and sagittal cross sections of the tumor segmentations (blue: necrosis; red: tumor; orange: edema). The second column shows the electric field distributions created using the clinical standard from the NovoTAL® planning software and the last two columns show the E-field intensity and E-field spread optimized results. Metrics: INT: average E-field magnitude in the tumor (higher is better); AF: coverage or “anti-focality” score (lower is better); E_brain_: average E-field magnitude in the whole brain; V_100_: Relative tumor volume over 100 V/m with respect to total tumor volume; V_225_: Relative tumor volume over 225 V/m with respect to total tumor volume.

**Table 1:** Comparison of the average electric field in the tumor (higher is better), the coverage score (anti-focality score, AF score, lower is better), and the Relative volume of the tumor area with respect to the total tumor volume in which the average electric field of the two stimulation channels exceeded values of 100 V/m between the current clinical standard and the optimized montages; Values in brackets are calculated over the whole brain. For every patient, the volume in mm³ of necrosis, tumor, and edema region is given in the first column.

|  | Clinical standard |  |  | Intensity optimized |  |  |  |  | Coverage optimized |  |  |  |  |
| --- | --- | --- | --- | --- | --- | --- | --- | --- | --- | --- | --- | --- | --- |
| Patient | Avg. E-field<br>in tumor<br>(V/m) | AF score | E-field<br>> 100 V/m<br>(whole<br>brain) | Avg. E-field in<br>tumor (V/m) |  | AF score |  | E-field<br>> 100 V/m<br>(whole brain) | Avg. E-field in<br>tumor (V/m) |  | AF score |  | E-field<br>> 100 V/m<br>(whole brain) |
| **L. occipital lobe**<br>$V_{necrosis} = 1,412 \text{ mm}^3$<br>$V_{tumor} = 4,682 \text{ mm}^3$<br>$V_{edema} = 34,312 \text{ mm}^3$ | 98.77 | 0.797 | 52.2%<br>(39.1%) | **121.31** | **+22.8%** | 0.742 | **-6.9%** | 74.5%<br>(32.4%) | 112.34 | **+13.7%** | **0.69** | **-13.4%** | 71.2%<br>(39.7%) |
| **R. frontal operculum**<br>$V_{necrosis} = 5,988 \text{ mm}^3$<br>$V_{tumor} = 27,067 \text{ mm}^3$<br>$V_{edema} = 84,487 \text{ mm}^3$ | 94.53 | 0.736 | 65.8%<br>(57.3%) | **116.79** | **+23.6%** | 0.640 | **-13.0%** | 80.3%<br>(44.9%) | 113.61 | **+20.2%** | **0.548** | **-25.5%** | 80.1%<br>(63.2%) |
| **R. temporal pole**<br>$V_{necrosis} = 0 \text{ mm}^3$<br>$V_{tumor} = 124 \text{ mm}^3$<br>$V_{edema} = 3323 \text{ mm}^3$ | 96.43 | 0.663 | 40.8%<br>(52.3%) | **129.25** | **+34.0%** | 0.575 | **-13.3%** | 99.8%<br>(49.8%) | 115.57 | **+19.6%** | **0.460** | **-30.6%** | 98.3%<br>(57.8%) |
| **R. frontal corpus callosum**<br>$V_{necrosis} = 5399 \text{ mm}^3$<br>$V_{tumor} = 19,599 \text{ mm}^3$<br>$V_{edema} = 73,201 \text{ mm}^3$ | 101.85 | 0.459 | 74.5%<br>(90%) | **122.11** | **+19.9%** | 0.322 | **-29.8%** | 79.4%<br>(82.5%) | 120.05 | **+17.9%** | **0.297** | **-35.3%** | 79.1%<br>(91.3%) |
| **R. inferior parietal lobe**<br>$V_{necrosis} = 13,553 \text{ mm}^3$<br>$V_{tumor} = 17,192 \text{ mm}^3$<br>$V_{edema} = 67,424 \text{ mm}^3$ | 83.05 | 0.840 | 29.1%<br>(57.3%) | **100.81** | **+21.4%** | 0.756 | **-10.0%** | 48.3%<br>(43.3%) | 98.79 | **+18.9%** | **0.673** | **-19.9%** | 47.1%<br>(59.9%) |

Additionally, the relative volume in the tumor area (with respect to the total tumor volume) in which the average electric field of the two stimulation channels exceeded values of 100 V/m was computed. The results are shown in Fig. 2 for the individual cases and are summarized in Table 1. The relative volumes were also calculated over the whole brain including tumor and are given in brackets in Table 1. The relative volumes exceeding a threshold of 100 V/m could be considerably increased by the optimization procedure compared to the standard placement.

In an additional analysis, the performance of the optimized configurations was compared against random placements of the electrode arrays. To this end, electric fields were computed for 400 randomized array configurations for each patient. Histograms of the average electric field and coverage score for one representative patient with a tumor in the right inferior parietal lobe are shown in Fig. 3. Similar histograms for the other patients are provided in the Supplementary Material (Fig. S2). These results indicate that the electrode configuration suggested by the NovoTAL® planning software outperforms the average scores from random placements, while the individual optimization algorithm yields further enhancements in performance metrics.

**Fig. 3:**
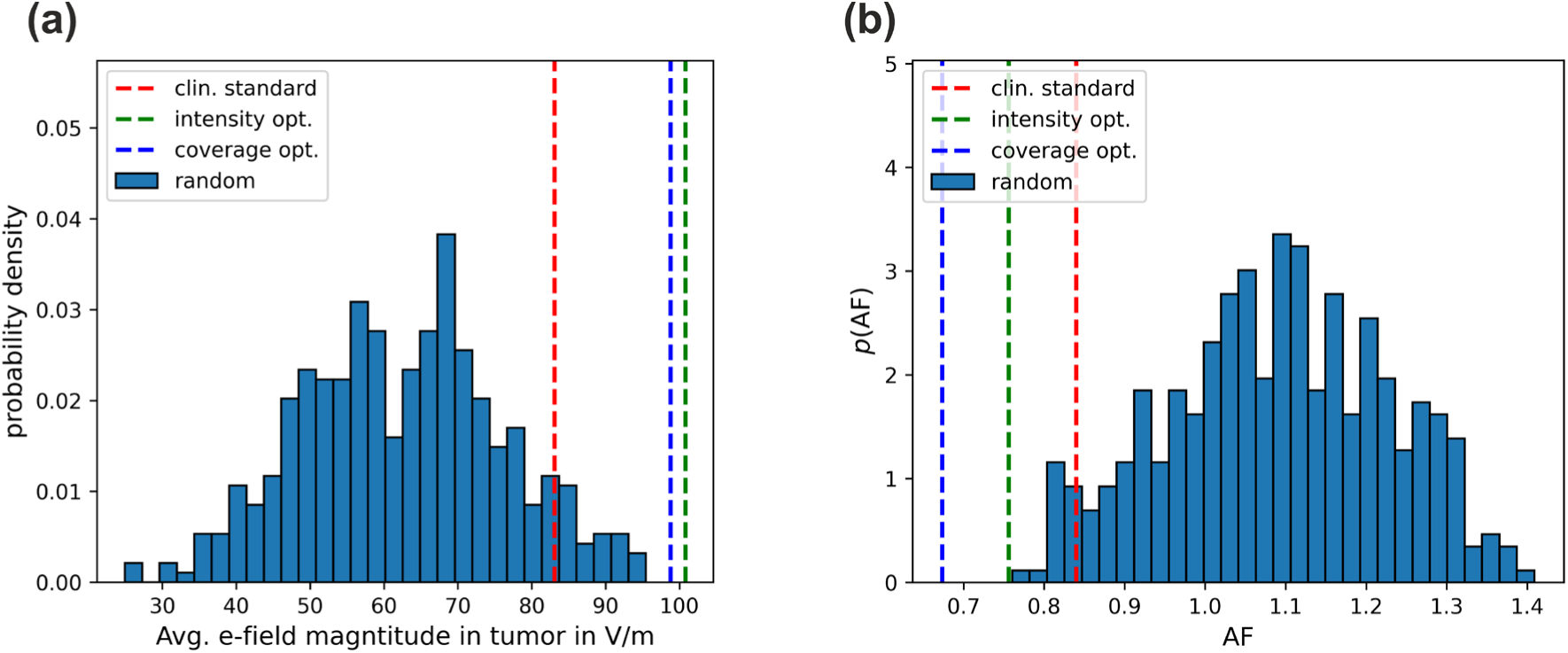
Histograms of the goal function values resulting from 200 random electrode placements for (a) E-field intensity optimization, showing the average electric field in the tumor (higher is better), and (b) optimization of the coverage score (anti-focality, AF, lower is better) from Weise et al. (2025). Results are shown for the patient with a tumor in the right inferior parietal lobe (last row in Fig. 2). The histograms of the remaining patients are shown in Fig. S2 in the supplemental material. The scores from the clinical standard using the NovoTAL® planning software are shown as red dashed lines.

The influence of tumor location and size on optimization outcomes was explored using artificial tumors modeled within the *ernie* head model. Fig. 4(a)-(c) illustrates the average electric fields achieved by the NovoTAL® software and the optimization algorithms for tumors as a function of tumor size as well as S-I and A-P tumor position. Notably, optimization substantially increased the absolute field strength for tumors near the surface (Fig. 4(a)). For A-P tumor locations (Fig. 4(b)), the optimization yielded a consistent improvement in the mean electric field strength by approximately 20% across all positions. Regarding tumor size (Fig. 4(c)), smaller tumors (≤15 mm) exhibited a higher benefit compared to larger ones (≥20 mm) from optimized electrode placements due to enhanced targeting accuracy.

**Fig. 4:**
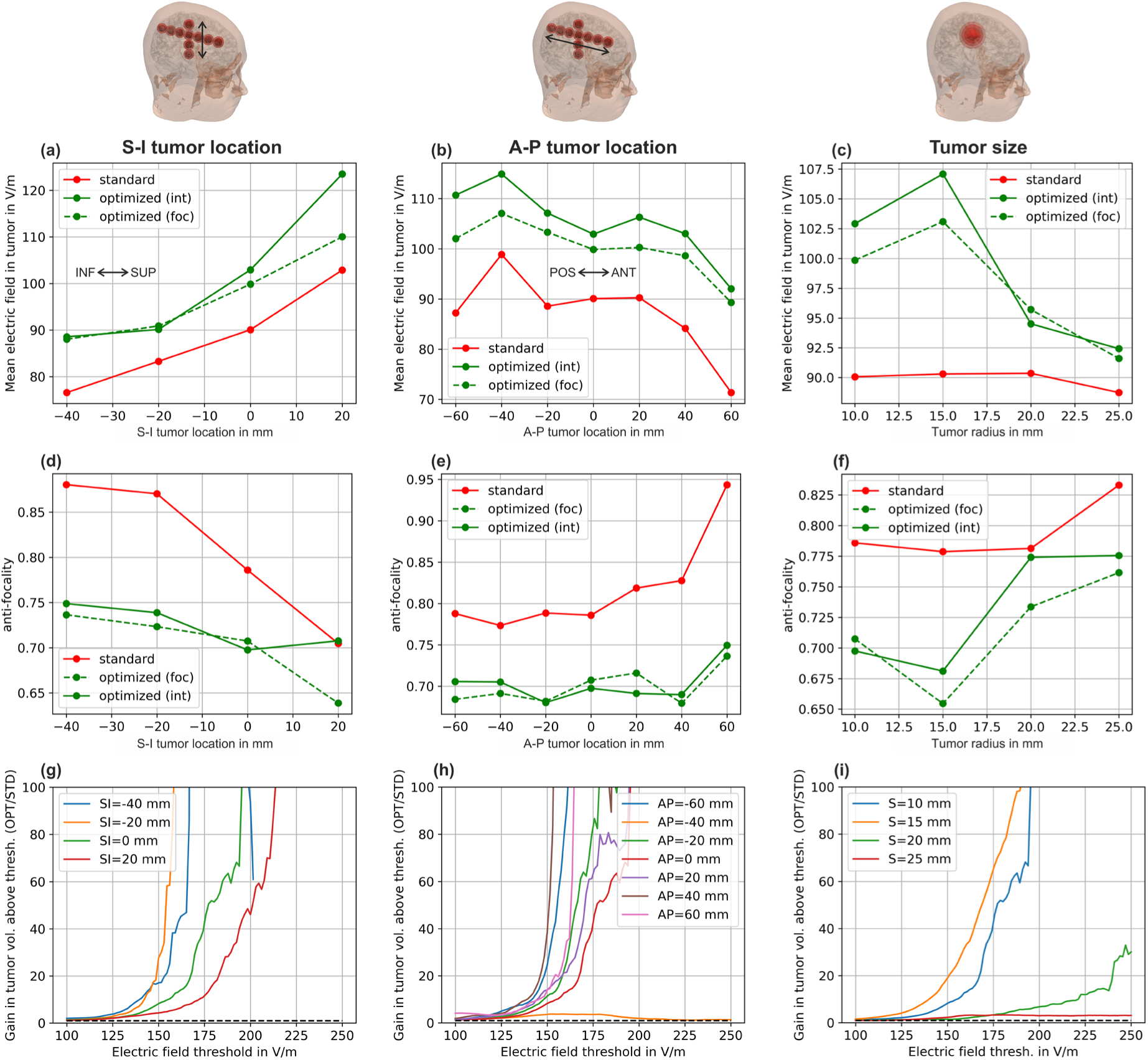
Comparison of electric field metrics between standard and optimized electrode positions for different tumor locations and sizes: (a)-(c) Average electric field in the tumor for S-I and A-P tumor locations, and tumor size, respectively (higher is better); (d)-(f) Coverage score (anti-focality, lower is better) in the brain for S-I and A-P tumor locations, and tumor size, respectively (lower is better); (g)-(i) Ratio of the relative volume of the tumor area with respect to the total tumor volume in which the average electric field of the two stimulation channels exceed a given electric field threshold (*x*-axis). The ratio is computed between intensity optimized and standard electrode positions. Values greater than 1 indicate the expected gain factor in relative tumor volume when using intensity optimization compared to the NovoTAL procedure.

Additionally, electrode configurations optimized for electric field coverage produced comparatively higher mean electric field strengths relative to standard configurations. Fig. 4(d)-(f) demonstrate that optimization focusing on field coverage resulted in significant improvements over standard electrode placements.

To further quantify these differences, partial tumor volumes exposed to electric fields exceeding specific thresholds were analyzed. The volume ratios between the optimized and standard configurations were calculated and are displayed in Fig. 4(g)-(i) for different tumor locations and sizes. Ratios exceeding one indicate increased field exposure, underscoring the better performance of optimized configurations in enhancing electric field coverage within the tumor region.

## Discussion

The findings of this study demonstrate further advancement in individually optimized TTFields electrode placements with the perspective of improving therapeutic outcomes. Using computational modeling, the approach consistently achieved better electric field distributions in the tumor region compared to the current clinical standard, as implemented by the NovoTAL® planning software. Specifically, the algorithm improved field coverage and enhanced the average field strength in the tumor by 18% to 34% across five representative patient cases. This improvement was further evident across various tumor locations, sizes, and configurations based on artificial head models, underscoring the robustness and adaptability of the optimization method. The results align well with recent work to improve the TTFields effects by personalizing the treatment^7^. For instance, Korshoej et al.^22^ demonstrated that patient-specific modeling improves field coverage and intensity in the glioblastoma. In that study, a manual search was used to find better electrode configurations compared to the current clinical standard, with the aim of demonstrating the possibility to improve beyond this standard. However, manual search is unfeasible in a clinical environment and also does not ensure the optimality of the solution. In contrast, numeric optimization offers a principled approach that yields optimal or close-to-optimal configurations, as we demonstrated by comparison with a large number of random configurations. The presented approach offers flexibility to adapt the optimization goal to the specific clinical use case – either maximizing the field in the tumor region or maintaining a balance between high field strengths in the tumor while also achieving broader field coverage. The latter option potentially enhances cytotoxicity in the main tumor region, while also targeting diffusely infiltrated brain areas. Collectively, the method enables flexible and objective optimization based on user-defined regions of interest and balanced clinical considerations.

The study highlights a clear influence of tumor size and location on electrode optimization. Smaller tumors and those located near the surface were particularly responsive to optimized placements, benefiting from more precise targeting. In the present simulation study, we observed that tumors with a radius smaller than 20 mm benefited most from the optimization. However, the targeting of the larger and more irregularly shaped tumors seen in some of the patient models was also clearly improved, highlighting the benefits of personalized optimization, including electric field simulations to address anatomical heterogeneity and complex tissue geometric profiles. This is currently not considered in standardized clinical dose-planning with NovoTAL®.

Despite promising results, certain limitations of our optimization method warrant consideration. First, while it demonstrated robust performance in simulated scenarios and patient-specific models, its efficacy in a clinical setting remains to be validated through randomized trials. Additionally, the dual optimization criteria—field intensity in the tumor target and coverage —provide a flexible framework. However, clinical and radiological criteria for balancing these factors, e.g. depending on the tumor type and distribution, still need to be developed.

It has been shown that TTFields’ efficacy depends strongly on TTFields dose^23,24^, which, according to Ballo et al.^9^, can be defined as the product of the TTFields average power loss density in the tumor bed *and* device usage (compliance). Power loss density and electric field magnitude are equivalent and related by the electrical conductivity of the tissue. Optimization offers the possibility of reducing the current amplitude to minimize side effects, which in turn could improve compliance and increase the TTFields dose in the long run.

Second, our study relied on static tumor models, which do not account for dynamic changes in tumor size or shape during treatment. Adjustment of array placement based on changes in tumor volume as seen on MRI would be important. Automatic optimizations, as introduced here, could be an important component to make such adjustments feasible in clinical practice. Third, manual application of electrode configurations, as described, may introduce user-dependent variability. Therefore, automating and systematically guiding this process could streamline clinical implementation of the optimized configurations. Fourth, other metrics in addition to the field intensity, including specific absorption rate (SAR) and current density, have been shown to impact TTFields efficacy^25,26^. SAR is a measure of energy deposition in tissue, influenced by factors such as tissue density — for example, the presence of cerebral edema around a tumor^27,28^. Lastly, at the cellular level, the field direction relative to the direction of cancer cell mitosis has been shown to play a role^1,6^. In pre-clinical research, increased TTFields efficacy was achieved using a combination of alternating orthogonal fields, yielding improved coverage of random mitotic directions. Previous studies have described alternative metrics of TTFields dose^17^ and related optimization methods^29^ to incorporate these factors in the evaluation of clinically applied electrode configurations. Future implementations of our algorithm may consider exploring these metrics in the objective function for more comprehensive optimization, although clinical validation is needed.

In addition to optimizing electrode placement, surgical approaches such as skull remodeling with cranial burr holes have been investigated to enhance Tumor Treating Fields (TTFields) intensity by mitigating the skull’s natural resistance to electrical currents^14,30–33^. However, recent results (Mikic et al. 2025, under review) indicate that such skull remodeling does not significantly improve treatment outcomes in recurrent glioblastoma, suggesting that dose enhancement may be less critical in this setting. Nevertheless, optimizing field distribution through surgical or planning-based methods could still hold relevance for patients with newly diagnosed glioblastoma, where TTFields efficacy may depend more strongly on achieving higher local field intensities.

In summary, this study underscores the potential of optimized electrode placement in enhancing the efficacy of TTFields therapy. By leveraging computational modeling and patient-specific anatomical data, the proposed optimization method significantly improved the electric field distribution compared to the current clinical standard. These improvements were consistent across various tumor types, sizes, and locations, highlighting the versatility and scalability of the approach. The findings emphasize the potential benefits of personalized TTFields treatment.

Looking forward, integrating adaptive and automated optimization systems could represent important next steps. Continuous monitoring of tumor changes for adjustment of electrode placement during treatment could further enhance the precision and efficacy of TTFields therapies. Validating the clinical benefits of these approaches in clinical trials will be crucial for translating them into clinical practice. Optimizing electrode placement represents an important advancement in TTFields therapy, offering a pathway to improved outcomes for patients with hard-to-treat cancers. By addressing the limitations of current clinical standards, this approach has the potential to increase the clinical efficacy of TTFields by a better personalization of the electrode positions.

## Code Availability

The approach is available in SimNIBS 4.5 (https://simnibs.github.io/simnibs/build/html/index.html). The optimizations can be carried out on a standard notebook with 16 GB RAM and take about 4-5h per patient to compute^16^.

## Data Availability

The datasets generated and analyzed during the current study are not publicly available due to privacy and ethical restrictions but may be shared on request. All shared data will be provided in anonymized form to protect patient confidentiality.

## Acknowledgements

The project was supported by the Danish Cancer Society (R322-A17630-B5570). AT was supported by the Lundbeck Foundation (grant R313-2019-622) and the National Institute of Health (grant R01MH128422). ARK was supported by the Independent Research Fund Denmark (9039-00307A) and Novocure, Ltd. KW and TRK were supported by BMBF grant 01GQ2201.

## Ethical statement

The study involved human participants, followed the Declaration of Helsinki, and was reviewed and approved by the Central Denmark Region Committee for Health Research Ethics (68928/1-10-72-214-19) and the Danish Data Protection Authority. Written consent was obtained from the trial participants for using non-identifiable imaging for future in silico studies.

## Credit Authorship Statement

KW, AT, TRK, and ARK formulated the overarching research goals and aims. KW conducted the analysis and ran the numerical simulations. AT reviewed the simulation code and results together with KW. ARK and NM provided the clinical data and ETW provided the electrode locations of NovoTAL®. NM provided the tumor segmentations and FC contributed to create the patient specific headmodels. ARK and ETW provided clinical interpretation of the results. TRK, ARK, AT supervised the study. KW wrote the initial draft and prepared the visualizations. ARK acquired the financial support. KW and TRK provided the computing resources. All authors critically reviewed the whole manuscript.

## References

1. Kirson ED, Gurvich Z, Schneiderman R, et al. Disruption of Cancer Cell Replication by Alternating Electric Fields. Cancer Res. 2004;64(9):3288–3295. doi:10.1158/0008-5472.CAN-04-0083

2. Mun EJ, Babiker HM, Weinberg U, Kirson ED, Von Hoff DD. Tumor-treating fields: A fourth modality in cancer treatment. Clinical Cancer Research.American Association for Cancer Research Inc. 2018;24(2):266–275. doi:10.1158/1078-0432.CCR-17-1117

3. Stupp R, Taillibert S, Kanner A, et al. Effect of Tumor-Treating Fields Plus Maintenance Temozolomide vs Maintenance Temozolomide Alone on Survival in Patients With Glioblastoma. JAMA. 2017;318(23):2306–2316. doi:10.1001/jama.2017.18718

4. Leal T, Kotecha R, Ramlau R, et al. Tumor Treating Fields therapy with standard systemic therapy versus standard systemic therapy alone in metastatic non-small-cell lung cancer following progression on or after platinum-based therapy (LUNAR): a randomised, open-label, pivotal phase 3 study. Lancet Oncol. 2023;24(9):1002–1017. doi:10.1016/S1470-2045(23)00344-3

5. Moser JC, Salvador E, Deniz K, et al. The mechanisms of action of Tumor Treating Fields. doi:10.1158/0008-5472.CAN-22-0887/3180540/can-22-0887.pdf

6. Wenger C, Miranda PC, Salvador R, et al. A Review on Tumor-Treating Fields (TTFields): Clinical Implications Inferred From Computational Modeling. IEEE Rev Biomed Eng. 2018;11:195–207. doi:10.1109/RBME.2017.2765282

7. Mikic N, Gentilal N, Cao F, et al. Tumor-treating fields dosimetry in glioblastoma: Insights into treatment planning, optimization, and dose–response relationships. Neurooncol Adv. 2024;6(1). doi:10.1093/noajnl/vdae032

8. Kirson ED, Dbalý V, Tovaryš F, et al. Alternating electric fields arrest cell proliferation in animal tumor models and human brain tumors. Proceedings of the National Academy of Sciences. 2007;104(24):10152–10157. doi:10.1073/pnas.0702916104

9. Ballo MT, Urman N, Lavy-Shahaf G, Grewal J, Bomzon Z, Toms S. Correlation of Tumor Treating Fields Dosimetry to Survival Outcomes in Newly Diagnosed Glioblastoma: A Large-Scale Numerical Simulation-Based Analysis of Data from the Phase 3 EF-14 Randomized Trial. International Journal of Radiation Oncology*Biology*Physics. 2019;104(5):1106–1113. doi:10.1016/j.ijrobp.2019.04.008

10. Glas M, Ballo MT, Bomzon Z, et al. The Impact of Tumor Treating Fields on Glioblastoma Progression Patterns. International Journal of Radiation Oncology*Biology*Physics. 2022;112(5):1269–1278. doi:10.1016/j.ijrobp.2021.12.152

11. Bomzon Z, Hershkovich HS, Urman N, et al. Using computational phantoms to improve delivery of Tumor Treating Fields (TTFields) to patients. In: 2016 38th Annual International Conference of the IEEE Engineering in Medicine and Biology Society (EMBC). IEEE; 2016:6461–6464. doi:10.1109/EMBC.2016.7592208

12. Wenger C, Salvador R, Basser PJ, Miranda PC. The electric field distribution in the brain during TTFields therapy and its dependence on tissue dielectric properties and anatomy: a computational study. Phys Med Biol. 2015;60(18):7339–7357. doi:10.1088/0031-9155/60/18/7339

13. Korshoej AR, Hansen FL, Thielscher A, von Oettingen GB, Sørensen JCH. Impact of tumor position, conductivity distribution and tissue homogeneity on the distribution of tumor treating fields in a human brain: A computer modeling study. PLoS One. 2017;12(6):e0179214. doi:10.1371/journal.pone.0179214

14. Korshoej AR, Saturnino GB, Rasmussen LK, von Oettingen G, Sørensen JCH, Thielscher A. Enhancing Predicted Efficacy of Tumor Treating Fields Therapy of Glioblastoma Using Targeted Surgical Craniectomy: A Computer Modeling Study. PLoS One. 2016;11(10):e0164051. doi:10.1371/journal.pone.0164051

15. Opitz A, Windhoff M, Heidemann RM, Turner R, Thielscher A. How the brain tissue shapes the electric field induced by transcranial magnetic stimulation. Neuroimage. 2011;58(3):849–859. doi:10.1016/j.neuroimage.2011.06.069

16. Weise K, Madsen KH, Worbs T, Knösche TR, Korshøj A, Thielscher A. A leadfield-free optimization framework for transcranially applied electric currents. Comput Biol Med. 2025;195:110648. doi:10.1016/j.compbiomed.2025.110648

17. Korshoej AR, Thielscher A. Estimating the Intensity and Anisotropy of Tumor Treating Fields Jsing Singular Value Decomposition. Towards a More Comprehensive Estimation of Anti-tumor Efficacy. In: 2018 40th Annual International Conference of the IEEE Engineering in Medicine and Biology Society (EMBC). IEEE; 2018:4897–4900. doi:10.1109/EMBC.2018.8513440

18. Thielscher A, Antunes A, Saturnino GB. Field modeling for transcranial magnetic stimulation: A useful tool to understand the physiological effects of TMS? In: 2015 37th Annual International Conference of the IEEE Engineering in Medicine and Biology Society (EMBC). IEEE; 2015:222–225. doi:10.1109/EMBC.2015.7318340

19. Puonti O, Van Leemput K, Saturnino GB, Siebner HR, Madsen KH, Thielscher A. Accurate and robust whole-head segmentation from magnetic resonance images for individualized head modeling. Neuroimage. 2020;219:117044. doi:10.1016/j.neuroimage.2020.117044

20. Weller M, van den Bent M, Preusser M, et al. EANO guidelines on the diagnosis and treatment of diffuse gliomas of adulthood. Nat Rev Clin Oncol. 2021;18(3):170–186. doi:10.1038/s41571-020-00447-z

21. Korshoej AR, Hansen FL, Mikic N, Thielscher A, von Oettingen GB, Sørensen JCH. EXTH-04. GUIDING PRINCIPLES FOR PREDICTING THE DISTRIBUTION OF TUMOR TREATING FIELDS IN A HUMAN BRAIN: A COMPUTER MODELING STUDY INVESTIGATING THE IMPACT OF TUMOR POSITION, CONDUCTIVITY DISTRIBUTION AND TISSUE HOMOGENEITY. Neuro Oncol. 2017;19(suppl_6):vi73–vi73. doi:10.1093/neuonc/nox168.300

22. Korshoej AR, Hansen FL, Mikic N, von Oettingen G, Sørensen JCH, Thielscher A. Importance of electrode position for the distribution of tumor treating fields (TTFields) in a human brain. Identification of effective layouts through systematic analysis of array positions for multiple tumor locations. PLoS One. 2018;13(8):e0201957. doi:10.1371/journal.pone.0201957

23. Ballo MT, Qualls KW, Michael LM, et al. Determinants of tumor treating field usage in patients with primary glioblastoma: A single institutional experience. Neurooncol Adv. 2022;4(1). doi:10.1093/noajnl/vdac150

24. Ballo MT, Conlon P, Lavy-Shahaf G, Kinzel A, Vymazal J, Rulseh AM. Association of Tumor Treating Fields (TTFields) therapy with survival in newly diagnosed glioblastoma: a systematic review and meta-analysis. J Neurooncol. 2023;164(1):1–9. doi:10.1007/s11060-023-04348-w

25. Lok E, San P, Hua V, Phung M, Wong ET. Analysis of physical characteristics of Tumor Treating Fields for human glioblastoma. Cancer Med. 2017;6(6):1286–1300. doi:10.1002/cam4.1095

26. Lok E, Clark M, Liang O, Malik T, Koo S, Wong ET. Modulation of Tumor-Treating Fields by Cerebral Edema from Brain Tumors. Adv Radiat Oncol. 2023;8(1):101046. doi:10.1016/j.adro.2022.101046

27. Panagopoulos DJ, Johansson O, Carlo GL. Evaluation of Specific Absorption Rate as a Dosimetric Quantity for Electromagnetic Fields Bioeffects. PLoS One. 2013;8(6):e62663. doi:10.1371/journal.pone.0062663

28. Wong ET, Lok E. Body Fluids Modulate Propagation of Tumor Treating Fields. Adv Radiat Oncol. 2024;9(1):101316. doi:10.1016/j.adro.2023.101316

29. Korshoej AR, Sørensen JChrH, von Oettingen G, Poulsen FR, Thielscher A. Optimization of tumor treating fields using singular value decomposition and minimization of field anisotropy. Phys Med Biol. 2019;64(4):04NT03. doi:10.1088/1361-6560/aafe54

30. Korshoej AR, Lukacova S, Lassen-Ramshad Y, et al. OptimalTTF-1: Enhancing tumor treating fields therapy with skull remodeling surgery. A clinical phase I trial in adult recurrent glioblastoma. Neurooncol Adv. 2020;2(1). doi:10.1093/noajnl/vdaa121

31. Korshoej AR, Mikic N, Hansen FL, Saturnino GB, Thielscher A, Bomzon Z. Enhancing Tumor Treating Fields Therapy with Skull-Remodeling Surgery. The Role of Finite Element Methods in Surgery Planning. In: 2019 41st Annual International Conference of the IEEE Engineering in Medicine and Biology Society (EMBC). IEEE; 2019:6995–6997. doi:10.1109/EMBC.2019.8856556

32. Cao F, Mikic N, Wong ET, Thielscher A, Korshoej AR. Guidelines for Burr Hole Surgery in Combination With Tumor Treating Fields for Glioblastoma: A Computational Study on Dose Optimization and Array Layout Planning. Front Hum Neurosci. 2022;16:909652. doi:10.3389/fnhum.2022.909652

33. Mikic N, Poulsen FR, Kristoffersen KB, et al. Study protocol for OptimalTTF-2: enhancing Tumor Treating Fields with skull remodeling surgery for first recurrence glioblastoma: a phase 2, multi-center, randomized, prospective, interventional trial. BMC Cancer. 2021;21(1):1010. doi:10.1186/s12885-021-08709-4

